# SETD2 Deficiency Promotes Inflammatory Bowel Disease via Oxidative Stress and FasL-induced Apoptosis

**DOI:** 10.1101/2022.11.10.516012

**Authors:** Yueduo Wang, Shenghai Shen, Li Li

## Abstract

SET-domain-containing 2 (SETD2) is known as the major trimethyltransferase that regulates the methylation of histone H3 lysine 36 (H3K36), which has been found frequently mutated in IBD samples. Although SETD2 deficiency has been proven to modulate oxidative stress, the specific mechanisms of SETD2 in IBD are still little known. To investigate the possible role of SETD2 in IBD, we generated and genotyped epithelium-specific deletion of *Setd2* (*Setd2*^Vil-KO^) progeny mice to establish mice IBD pathology model and then studied molecular expression differences at mRNA and protein levels in the intestinal epithelial tissue of *Setd2*^Vil-KO^ mice. Compared with *Setd2*^F/F^ mice, the tissue of *Setd2*^Vil-KO^ mice showed increases in oxidative stress and cell apoptosis level via RT-qPCR, while TUNEL assay and immunofluorescence showed significantly enhanced apoptotic signaling and FasL expression. Together, our findings highlight that SETD2 regulates oxidative stress and FasL-induced apoptosis in *Setd2*^Vil-KO^ mice intestinal epithelial cells through epigenetics mechanisms, which plays an essential role in IBD pathogenesis. Therefore, our study provides new insights into the prevention and therapy of IBD from the perspective of epigenetic regulation.

## 1. Introduction

Inflammatory bowel disease (IBD) is a chronic and recurrent inflammatory disorder of the mucosal barrier that can be sub-classified into Crohns disease (CD) and ulcerative colitis (UC) [^1,2^]. The mucosal barrier is composed of a variety of cells, including intestinal epithelial cells (IECs), one of the most active self-renewal tissue cells in adult mammals, which adhere to each other via tight junctions and are essential for maintaining intestinal epithelium homeostasis [^3^]. The integrality of the mucosal barrier is usually affected by multiple factors, such as the invasion of pathogenic microbes, apoptosis, and oxidative stress[^1,3–5^]. Oxidative stress is one of the common factors that initiation by exceeded reactive oxygen species (ROS) accumulation, which induces cell apoptosis, and results in mucosal barrier disorder, inflammation, and intestinal epithelium homeostasis damage [^3,4^]. In addition, the continuation of tissue inflammation would cause tissue repair obstacles and tumorigenesis. Hence, long-term IBD is more likely to result in colorectal cancer (CRC) [^3^], and therefore it is imperative to prevent IBD development and deterioration, which includes transforming into CRC.

The term “epigenetics” was proposed to describe the DNA templating events based on chromatin, such as histone modifications and nucleosome remodeling [^6^]. Epigenetics regulators play a critical role in many biological processes and diseases, including IBD development [^3^]. In relative studies, EZH2, an epigenetics regulator that inactivates the trimethylation modification of histone H3 lysine 27, was found to promote inflammatory response and cell apoptosis due to its exceed expression, which contributes to mucosal barrier disorder and IBD development [^1,7^]. Moreover, BRG1 is an essential regulator for autophagy gene transcription in the colon, which attenuates colitis and tumorigenesis through autophagy process-dependent ROS isolation [^4^]. In previous studies, SETD2 was also found as a key epigenetics regulator for the trimethylation of H3K36, whose mutation rate reached about 17% in IBD samples [^3,7^].

SETD2 is the central regulator that alters the trimethylation states of lysine 36 on histone 3 (H3K36me3) in mammalian cells, which plays a critical role in maintaining genomic integrity and stability such as transcriptional regulation, DNA damage repair, chromosome segregation, and alternative splicing [^8,9^]. The mutation of SETD2 in actively transcribed regions could serve as a potential multifunctional histone biomarker due to its unique features [^3,9^]. Therefore, SETD2 has been extensively studied in various biological processes and diseases recently, such as prostate cancer metastasis [^7^], hepatocarcinoma [^10^], as well as the development of pancreatic cancer and initiation of normal lymphocytic development [^3^]. However, the specific functions of SETD2 in IBD pathogenesis remain little known.

To investigate a possible role of SETD2 in IBD, we established *Setd2*^Vil-KO^ mice and then induced with DSS to generate colonic inflammation. RT-qPCR, TUNEL assay, and immunofluorescence were performed to detect molecular expression differences in the intestinal epithelial tissue of *Setd2*^Vil-KO^ mice, and our study showed that SETD2 deficiency leads to FasL-induced apoptosis and increased oxidative stress levels in *Setd2*^Vil-KO^ mice.

## 2. Materials and methods

### 2.1. IBD Mice pathology model establishment

*Setd2*-flox (*Setd2*^F/F^) mice and Villin-Cre transgenic mice were purchased from Shanghai Biomodel Organism Co., bred and maintained under pathogen-free conditions and an accredited animal facility. *Setd2*^F/F^ mice were crossed with Villin-Cre mice to establish *Setd2*^Vil-KO^ mice, and both *Setd2*^F/F^ mice and *Setd2*^Vil-KO^ mice were then treated with inducing agent dextran sodium sulfate (DSS) to induce IBD. Both *Setd2*^F/F^ and *Setd2*^Vil-KO^ mice were given 3% DSS daily for 7 days, with drinking water regularly followed for 5 days. Subsequently, 2% DSS was given for 5 days, with drinking water regularly followed for 5 days. All experiments were conducted on C57BL/6J mice with littermates as controls, and all the procedure has been approved by the Ethics Review Board of Renji hospital.

### 2.2. DNA extraction from mouse tail for genotyping

Each mouse tail tip was cut and put into one eppendorf tube separately, added with 100 μL to 150 μL reagent A (25mM NaOH/0.2mM EDTA, 5×stock solution that can be kept at room temperature) to each eppendorf tube, boiled at 95 to 98°C for 1 hour and cool down at room temperature. Then add an equal volume of reagent B (40mM Tris HCl, pH=5.5) to each Eppendorf tube and mix well, centrifuge at full speed for 5 minutes and 1 μL to 2 μL supernatant were taken for PCR. Nucleic acid electrophoresis was performed to genotype.

### 2.3. RNA Isolation and Real-Time PCR

Total RNA of the intestinal epithelium tissue of *Setd2*^Vil-KO^ mice was extracted with TRIzol kit (Invitrogen), following the manufacturers instructions. 2 g of total RNA and SuperScript II (Invitrogen) were used to synthesize first-strand cDNA, then the reaction was performed using SYBR Green Universal Master Mix reagent (Roche). Serial dilutions of total RNA were used to generate standard curves, and all mRNA levels were normalized against β-actin mRNA. All samples were subjected to three parallel experimental replicates.

### 2.4. The TUNEL assay and immunofluorescence

The intestinal epithelium tissue of *Setd2*^Vil-KO^ mice was handled by using the swiss roll technique, 4% paraformaldehyde was then used as a fixative overnight. Treated tissue was embedded in paraffin, and cut into 5-mm sections, and paraffin sections were then rehydrated and heated to induce antigen retrieval. Blinded scoring was performed on H&E-stained sections, and samples from all sections of the colon were examined by pathologists. The apoptotic signaling in the intestinal epithelial tissues of *Setd2*^Vil-KO^ mice was detected by TUNEL staining, using the TUNEL Assay Kit (Fluorescence, 488 nm) #25879 CST. Primary antibodies used for immunofluorescence were anti-fasL (sc-19681,200ug/ml). Both TUNEL assay and immunofluorescence were observed under the fluorescence microscope.

### 2.5. Statistical Analysis

Student’s t-test was used to analyze the data in RT-qPCR, and results were presented as means ± standard errors of the mean (SEM). All data sets were analyzed with the GraphPad Prism. A p < 0.05 is considered significant.

## 3. Results

### 3.1. SETD2 deficiency altered antioxidative and apoptosis gene expression in Setd2Vil-KO mice

To investigate the possible role of SETD2 in IBD, we collected RNA-seq data from the study of Min L et al. [4]. In RNA-seq data, 644 up-regulated genes and 605 down-regulated genes (fold change > 1.25) were found in a total of 18168 differentially expressed genes, while gene ontology analysis (GO) indicated that significant enrichment for genes related to apoptosis regulation and inflammatory response, as well as antioxidative genes. To identify the major differentially expressed genes, we performed a secondary analysis based on the maximum of log2FC absolute value, and four major differentially expressed genes were finally identified (Table. 1). Among which redox genes cytochrome b-245 heavy chain (C*ybb*) and lactoperoxidase (*Lpo*) were showed down-regulated, while two other genes associated with cell apoptosis, Fas ligand (*FasL*) and tumor necrosis factor receptor superfamily member 8 (*Tnfrsf8*) expression were detected enhanced versus *Setd2*^F/F^ mice. To validate these results, we established IBD mice pathology model (Fig.1) and performed RT-qPCR, which consistent with the results in RNA-seq (Fig.2). *Cybb* and *Lpo* expression were decreased about thereefold and fourfold respectively, while *FasL* and *Tnfrsf8* expression increased about threefold and 0.7 times respectively. Together, these results emphasize that SETD2 deficiency is responsible for the down-regulation antioxidative genes C*ybb* and *Lpo*, as well as the up-regulation of apoptosis genes *FasL* and *Tnfrsf8* at the mRNA level.

**Table. 1.**
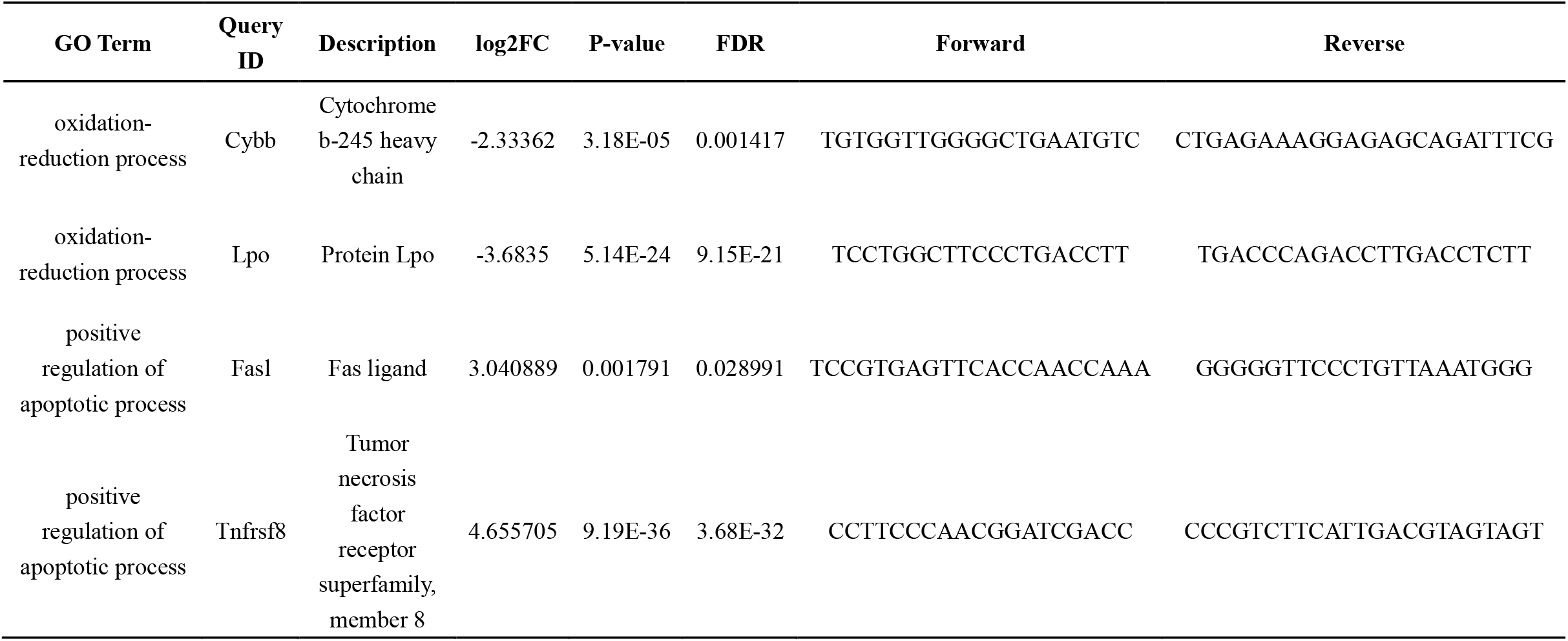
RNA-seq analysis of associated gene expression differences in *Setd2*^Vil-KO^ mice. Four major differentially expressed genes with P-value<0.05 and FDR <0.05 were identified with maximum log2FC absolute value. These genes were significantly enriched with the oxidation-reduction and positive regulation of the apoptotic process due to the GO term.

**Fig. 1.**
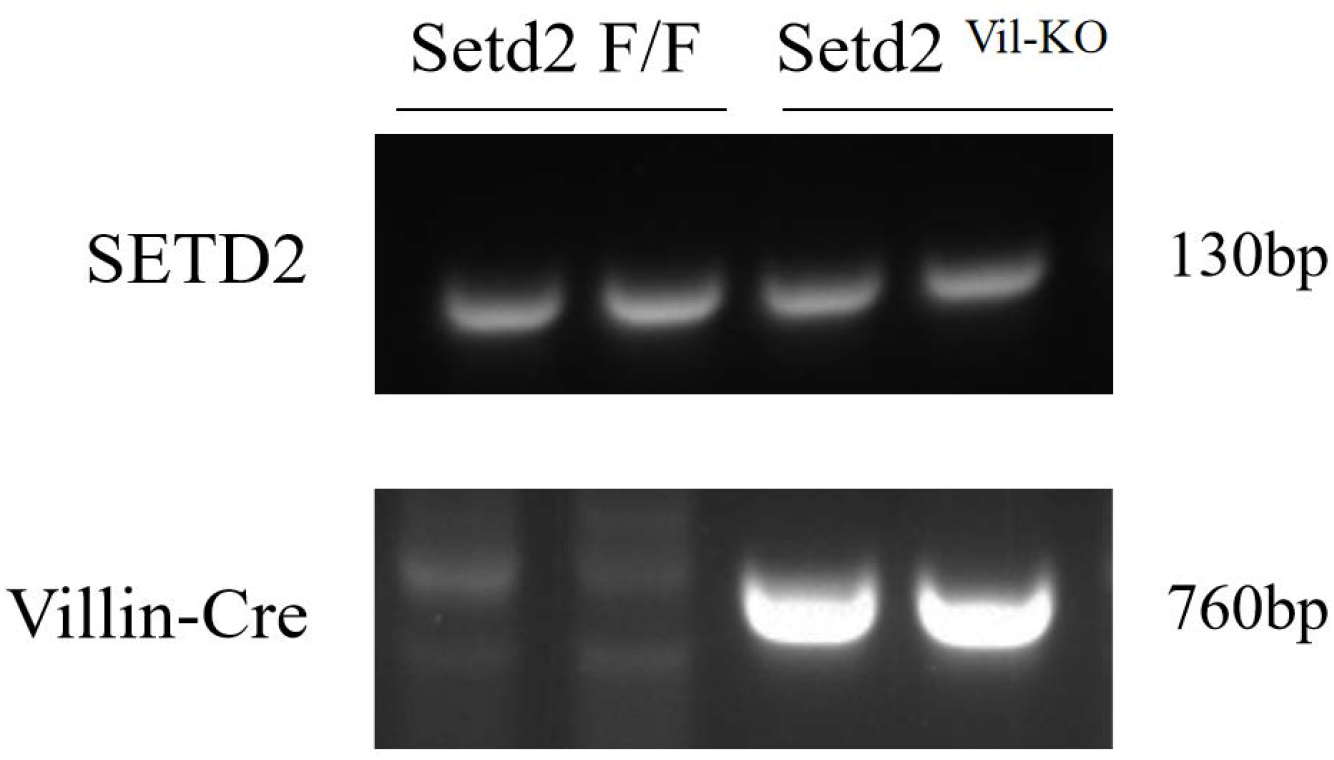
The genotyping of Setd2^Vil-KO^ mice model.

**Fig. 2.**
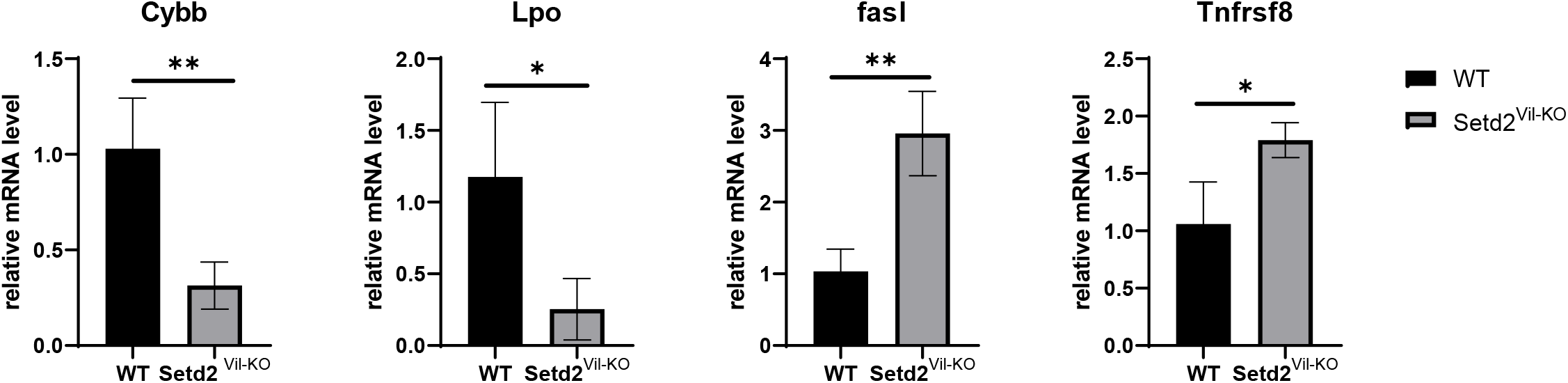
RT-qPCR analysis of associated gene expression differences in Setd2Vil-KO mice. Students t-test was used to determine the statistical significance of the mean±S.E.M. (n = 3 per genotype, *p < 0.05, **p < 0.01. N.S., Not Significant.)

### 3.2. SETD2 deficiency enhanced apoptotic signaling in Setd2Vil-KO mice

Due to the significant enhancement of *FasL* and *Tnfrsf8* expression at mRNA level in RNA-seq and RT-qPCR, apoptosis might therefore be one factor that is highly related to IBD development. To further evaluate the relevance of apoptosis in IBD, we then performed TUNEL assays to detect apoptotic signaling in *Setd2*^Vil-KO^ mice intestinal epithelium tissue. Corresponds with our results in RNA-seq and RT-qPCR, apoptotic signaling was detected to be significantly up-regulated in the intestinal epithelium tissue of *Setd2*^*Vil-KO*^ mice (**Fig.3**), which represents that cell apoptosis process in the intestinal epithelium tissue of *Setd2*^Vil-KO^ mice was increased.

**Fig. 3.**
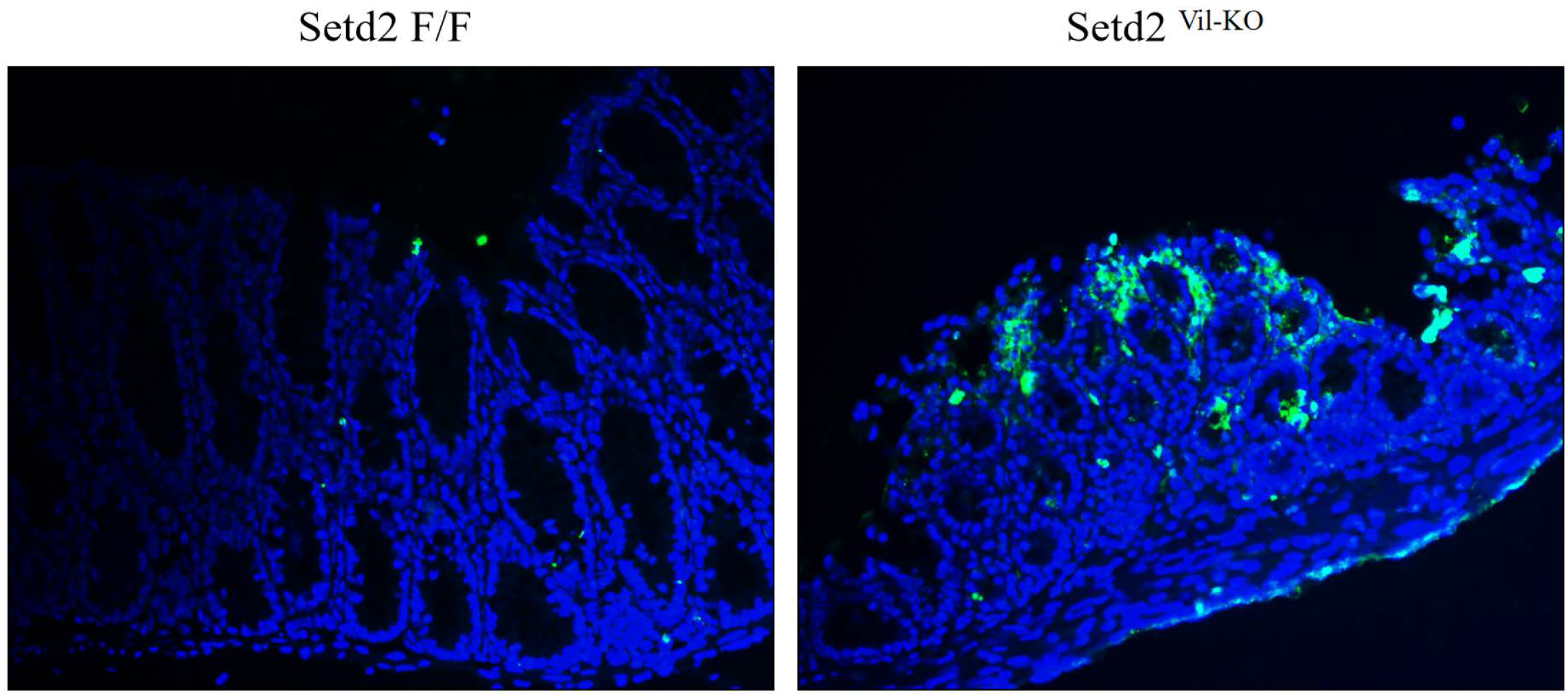
TUNEL assay of Setd2^Vil-KO^ mice intestinal epithelial tissues. Green fluorescence represents apoptotic signaling. (Scale Bars: 50 μm.)

### 3.3. SETD2 deficiency leads to FasL-apoptosis in Setd2Vil-KO mice

Towards a molecular basis of cell apoptosis in the intestinal epithelium tissue of *Setd2* deletion mice, we subsequently performed immunofluorescence to detect FasL expression at the translation level. Consistent with our findings in the TUNEL assay, FasL was found to be significantly up-regulated at the translation level in the intestinal epithelium tissue of *Setd2*^Vil-KO^ mice (Fig.4). Collectively, FasL was found to be up-regulated at both mRNA (approximately 3 times) and protein levels in the intestinal epithelium tissue of *Setd2*^Vil-KO^ mice. Thus, we demonstrated that SETD2 deficiency leads to FasL-apoptosis in *Setd2*^Vil-KO^ mice intestinal epithelial tissues.

**Fig. 4.**
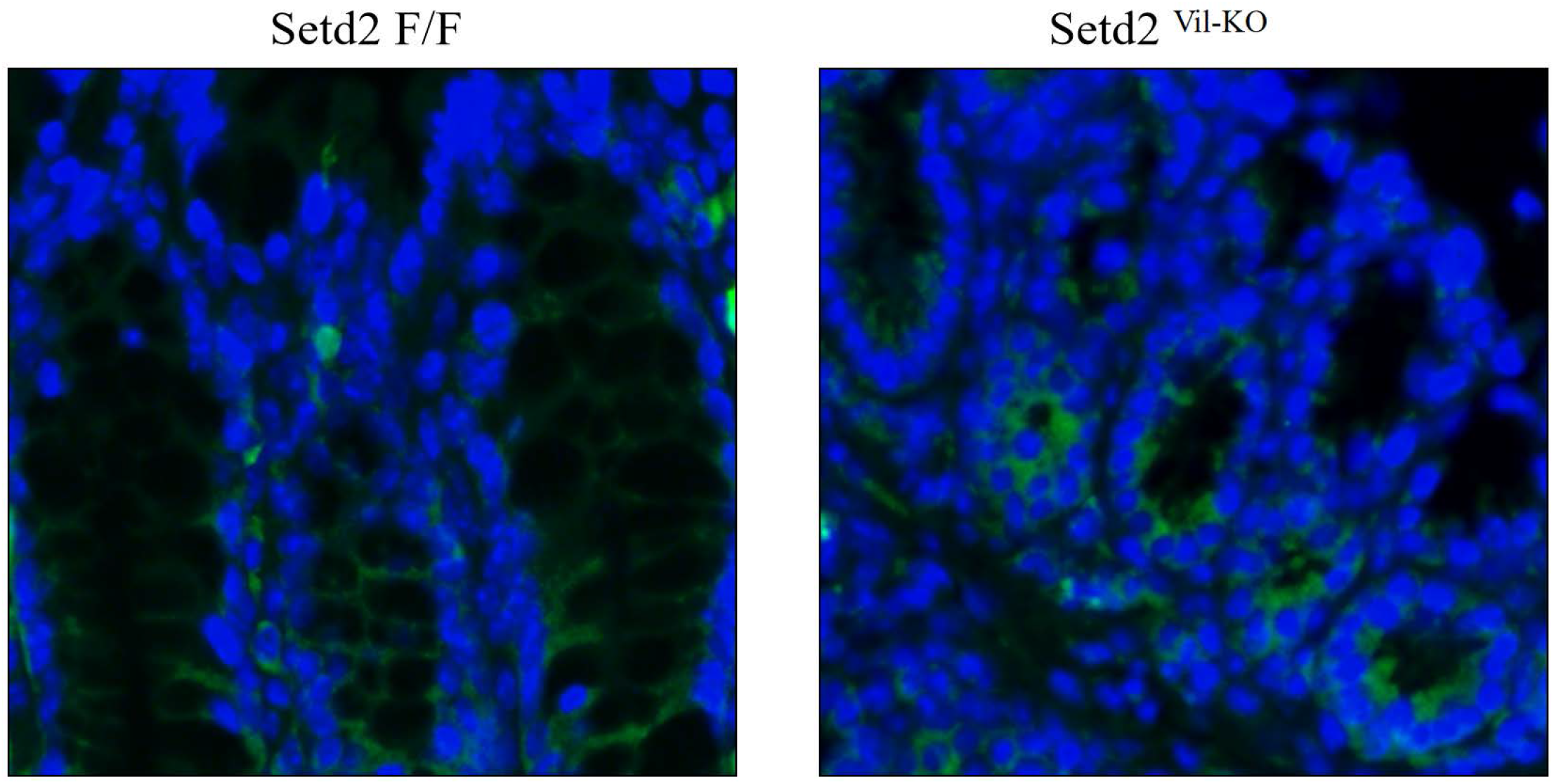
The immunofluorescence of Setd2^Vil-KO^ mice intestinal epithelial tissues. Green fluorescence represents FasL protein. (Scale Bars: 100 μm.)

## 4. Discussion

IBD is a chronic inflammatory disorder of the mucosal barrier with multiple triggers, among which epigenetics regulations such as DNA methylation, histone modifications, and nucleosome remodeling were found to play an essential role in IBD development. As a key epigenetics regulator, SETD2 is known as the major trimethyltransferase of histone H3K36, which was detected frequently mutated in IBD patients, and has been proven to play a critical role in many diseases. However, the specific functions of SETD2 in IBD development remain unclear. To investigate the possible role of SETD2 in IBD pathogenesis, *Setd2* epithelium-specific deletion mice (*Setd2*^Vil-KO^ mice) were established, and DSS was then treated to generate IBD. We demonstrated that the oxidative stress level and apoptotic cell signaling in *Setd2*^Vil-KO^ intestinal epithelium were enhanced compared with *Setd2*^F/F^ mice. Our findings showed that SETD2 deficiency leads to FasL-induced apoptosis and promotes oxidative stress in *Setd2*^Vil-KO^ mice, which supports the findings in other studies that SETD2 is frequently mutated in IBD patients. Thus, we provided evidence that SETD2 plays a critical role in IBD pathogenesis.

This study reveals a mechanism that SETD2 deficiency leads to FasL-apoptosis and promotes oxidative stress in IBD mice. It is well known that ROS contributes to many disease developments, such as cardiovascular disease, inflammation, and cancer [^11,12^]. We found that the expression of antioxidative genes *Cybb* and *Lpo* were significantly down-regulated about 3 times and 4 times, respectively. CYBB is a primary component of the phagocyte NADPH oxidase (NOX) system [^13^]. The disorder of the NOX system leads to granulomatous disease (GCD), which is characterized by fatal bacterial, and fungal infections and inflammation, and about one-third of GCD patients develop into including IBD [^13,14^]. Moreover, lactoperoxidase (*Lpo*) is a role in scavenging ROS [^15^], and hence the down-regulation of *Cybb* and *Lpo* indicates that the antioxidant capacity was reduced, and hence the oxidative stress level was promoted in the intestinal epithelium of *Setd2*^Vil-KO^ mice. This finding is also supported by the study of Min L et al., among which 906 antioxidative genes in total, including *Prdx3, Prdx6, Gclm*, and *Srxn1*, were significantly suppressed at mRNA expression levels after the ablation of SETD2 in the intestinal epithelium tissue of *Setd2*^Vil-KO^ mice [^3^]. Therefore, SETD2 deficiency is mainly responsible for the enhancement of oxidative stress levels. Cell apoptosis is critical for eliminating damaged, senescent, or useless cells without causing microenvironment damage and inflammation [^16^], which has been found to accelerate in IBD patients [^17^]. In this research, significant up-regulation of *FasL* and *Tnfrsf8* was detected, among which *FasL* expression was increased 3 times approximately, while *Tnfrsf8* expression was enhanced about 0.7 times. FasL is a key death factor in the tumor necrosis factor (TNF) family for receptor-triggered programmed apoptosis bounds to Fas [^18^]. Normally, FasL is low expressed in IECs, [^19^]. However, a high expression level of FasL mRNA was detected in UC patients while maintained at a normal level in CD patients [^20,21^], consistent with our results. Furthermore, we also found apoptotic signaling and FasL translation level were both increased in *Setd2*^Vil-KO^ mice intestinal epithelium, which is a shred of solid evidence that FasL was up-regulated at both mRNA and protein levels. Therefore, our findings reveal for the first time that SETD2 deficiency leads to FasL-apoptosis and uncover part of the potential association of epigenetics regulation with apoptosis in the pathogenesis of IBD.

It is worth mentioning that the tumor necrosis factor (TNF) family appears to play an essential role in SETD2 loss IBD mice. Notably, both Tnfrsf8 (CD30) and FasL belong to the TNF superfamily [^18,22^], and CD30 signaling was found to up-regulate *Fas* gene expression [^23^]. However, little is known about the specific functions of TNF family members in SETD2-mutated IBD patients. Considering that apoptosis is a complicated biological process, which usually with multiple types and triggers. Therefore, it remains to be determined whether SETD2 mutation leads to other types of cell apoptosis in IBD development, and the mechanism of epigenetic regulation triggered by SETD2 loss in IBD has also not been fully defined yet.

## 5. Conclusion

SETD2, as an important enforcer of the epigenetic regulator, is widely involved in various diseases, such as prostate cancer metastasis, hepatocarcinoma, and the initiation of normal lymphocytic development. In this study, four genes were identified as major differential genes, among which antioxidative genes *Cybb* and *Lpo* were significantly down-regulated about 3 times and 4 times, respectively, while apoptosis genes *FasL* and *Tnfrsf8* were up-regulated about 3 times and 0.7 times, respectively. These results indicate that compared with wild-type mice, the intestinal epithelial tissues of *Setd2*^Vil-KO^ mice undergo reduced antioxidant capacity and enhanced apoptosis, and hence SETD2 deficiency leads to the enhanced oxidative stress level. Furthermore, increased apoptotic signaling and up-regulated FasL expression at both mRNA and protein levels were found in the intestinal epithelial tissues of *Setd2*^Vil-KO^ mice, which suggests that SETD2 deficiency leads to FasL-apoptosis. Overall, this study first established the relationship between *Setd2* deletion and cell apoptosis in the context of IBD. However, our study only focused on FasL-apoptosis, while other types of cell apoptosis led by SETD2 mutation in IBD development are still little known. Moreover, the mechanism of epigenetic regulation triggered by SETD2 loss in IBD is still poorly understood. Therefore, the potential relationship between epigenetic regulation and apoptosis can be used as an entry point for future studies on the pathogenesis of IBD, thereby providing deeper insights into the pathogenesis, treatment, and prevention of IBD from the perspective of molecular pathology. Considering that SETD2 mutation account for a higher risk of developing into CRC in IBD patients (about 17%), Our results will provide theoretical support and basis at the molecular level for the mechanism, therapy, and prevention of such diseases.

## Author contributions

Conceptualization, Y.W(leading). and S.S(support).; Formal analysis, S.S.; Funding acquisition, L.L.; Investigation, Y.W. and S.S; Methodology, Y.W.; Project administration, L.L.; Resources, L.L.; Supervision, L.L.; Validation, Y.W.; Visualization, Y.W. and S.S; Writing—original draft, Y.W.; Writing—review & editing, Y.W. and S.S. All authors have read and agreed to the published version of the manuscript.

## Data accessibility

All data are available from the authors upon reasonable request. RNA-Seq and ChIP-Seq raw data have been deposited in the Gene Expression Omnibus (GEO) under accession number GEO: GSE 151968.

